# Insights from Genetic Studies: SNP Analyses Confirm White Clover Naturalization in Brazil

**DOI:** 10.1101/2024.02.09.579627

**Authors:** Amanda S Alencar, Yoshishisa Suyama, Daiki Takahashi, Vidal de F Mansano, Catarina da F Lira

## Abstract

White clover (*Trifolium repens*) is a stoloniferous legume herb native to Eurasia, which had been introduced and spread globally. In Brazil, it was introduced as forage crop. While previous studies focused mainly on its agricultural benefits, much remains unclear about its territorial dynamics, introduction process and potential threats in Brazil. This study aims to estimate the genetic diversity of naturalized white clover populations in Brazil and assess the influence of cultivars into these populations’ diversity. Through MIG-Seq analysis, 1097 SNPs show that Brazilian populations have 94% within-population variation. Additionally, two mountainous areas clustered together, while rural and urban areas formed a second cluster. Cultivars are less diverse and have 27% of their genetic variability between them. We found that some populations admixture with cultivated varieties, while more isolated mountainous populations were singular in their genetic background. We can conclude that it is possible that parts of the populations are originally native, brought during European immigration, while others appear to have similar cultivar ancestry, indicating possible biological escapes from cultivars into naturalized populations. Considering ecological data and our genetic findings, it is confirmed that white clover is indeed naturalized in Brazil.

## Introduction

Genetic variability, the intrinsic adaptive potential of organisms (Aravanopoulos, 2011), is influenced by various factors and evolutionary forces. Among these, the process of colonizing new territories plays a crucial role in determining the initial genetic pool of populations. For instance, the occurrence of single or multiple species’ introduction events influences the frequency of acquiring new genetic variability during the colonization process. The introduction of variability is dependent on the origin and diversity of genotypes involved in the process directly impacting the strength of the founder effect (Reed et al. 2003; Blackburn et al. 2016).

The capacity of a species to disperse also influences its genetic variability. Species with high dispersive capacity tend to exhibit greater gene flow and, consequently, higher genetic variability. However, habitat fragmentation and anthropogenic activities can accentuate the loss of variability through genetic drift events, such as bottlenecks (Johnson and Munshi-South 2017). Plant species are markedly affected by fragmentation. Unable to migrate between habitats rapidly, these species may also experience disruptions in pollinator-plant interactions and seed dispersal patterns (Leimu et al. 2015), rendering them vulnerable organisms in need of conservation actions.

The Global Strategy for Plant Conservation (GSPC) is one of the initiatives that Brazil and other countries have joined to preserve plant biodiversity. The objective is for signatories to allocate human and financial resources to enhance knowledge, conservation and sustainable use of plant biodiversity at local and global scales (CDB, 2011). Among the 16 established goals, part of goal four aims to have “70 percent of the genetic diversity of crops and other important plant species of great socioeconomic value conserved”. To attain this goal, promoting genetic studies is essential. Ottewell and collaborators (2015) have already highlighted that despite its recognized importance in conservation plans alongside ecological data, genetic diversity preservation does not yet receive the necessary emphasis.

*Trifolium repens* (Fabaceae), known as white clover, is a stoloniferous perennial legume native to subtropical and temperate climates (Sabudak & Guler 2009). It exhibits both vegetative and sexual reproduction through pollinators from the genera *Apis* and *Bombus* playing a crucial role (Larson et al. 2014), since white clover is self-incompatible (Bissuel-Belaygu et al. 2002). Its seeds can be ingested by birds and ruminants, and dispersed over long distances after passing through the gastrointestinal tract (Yamada and Kawaguchi 1972). Despite its Eurasian origins, white clover is found on every continent (Griffiths et al. 2019). In Brazil, its introduction was for forage purposes (Leite et al. 2018), but no information regarding its origin is available.

White clover is one of the most used temperate legumes by livestock farmers in the southern region of Brazil (Lorenzi 2000; Rosa Neto et al. 2014). Typically sown in intercrop (Dall’Agnol et al. 1982), it has high biomass production, nutritional value and enhances the palatability of pastures for livestock. Additionally, it is recommended for overseeding and soil recovery due to its nitrogen fixation capacity (Caradus et al. 1996; Hoyos-Villegas et al. 2019). Since the 1920s, more than 320 species of cultivars of the species have been developed worldwide (Caradus and Wood-field 1997; Hoyos-Villegas et al. 2019). Natural populations of white clover are distributed in the Southeast and South regions of Brazil, accounting for 80% of the recorded occurrences (Specieslink 2020). They inhabit diverse environments, particularly Highland fields (Safford 1999), clean fields (Trifolium in Flora do Brasil 2020) and areas of association with grasses in urban and rural environments (Alencar personal observation).

Hargreaves et al. (2010) demonstrated the occurrence of admixture and gene flow between native and cultivated white clover populations across insular and continental regions of the United Kingdom. This suggests a tendency towards increased genetic similarity between these populations, albeit without complete homogenization of the genetic pool. In clonal plants such as white clover, genetic diversity can mitigate genetic monomorphisms (Verhieven and Preite 2014; González et al. 2016). These studies underscore the importance of incorporating common species with gene flow with cultivars into conservation plans, while also integrating genetic data into the analysis.

Single nucleotide polymorphisms (SNPs) are specific polymorphisms throughout the genome. The SNPs can be accessed through whole-genome sequencing techniques. Multiplexed ISSR genotyping by sequencing (MIG-seq), a high-throughput genotyping technique that utilizes next-generation sequencing (NGS), is employed to analyze genetic diversity and population structure (Suyama and Matsuki, 2015; Suyama et al 2022). MIG-seq offers cost-effective genotyping of large sample sizes without requiring extensive optimization. Its broad applications span ecology, evolution, conservation biology, breeding, and crop improvement (Sakaba et al. 2023, Suetsugu et al. 2023a, Suetsugu et al. 2023b, Takahashi and Suyama 2023). MIG-seq facilitates marker identification, population structure analysis, and evolutionary studies, making it a versatile and efficient tool for understanding genetic variation and its biological implications.

Previous studies on white clover in Brazil have primarily been focused on applied agronomic issues related to the species, such as seed production, biomass production for foraging, and related topics. Consequently, our understanding of the dynamics of white clover’s territorial conquest and its establishment in Brazil remains limited. These knowledge gaps hinder the accurate classification of threats to the species and allow for blind spots in controlling its colonization behavior and potential management.

In light of objectives such as the GSPC and the global significance of the species, it is crucial to invest resources in filling these knowledge gaps to advance biodiversity conservation. Therefore, this study aims to provide a genetic overview of naturalized white clover populations in Brazil and comprehend how cultivars contribute to the genetic structure of the species within the territory.

## Materials and methods

### Brazilian plant material

From October 2020 to January 2021, a total of 30 white clover individuals were randomly collected (Supplemetary material S1) from five locations in the cities of Gramado (RS; RSG), Caxias do Sul (RS; RSCS), São Bento do Sul (SC; SCBS), Lages (SC; SCL), Curitiba (PR; PRC) (Figure 1A), Teresópolis (RJ; RJT) and within the limits of the Itatiaia National Park (which comprehends three different cities, but here will be referred as RJI) (Figure 1B). Fully expanded leaves were carefully collected, individually stored in cloth sachets labeled with unique sample codes, and dried in silica gel until utilized for molecular analysis. To minimize the possibility of collecting clonal replicates, the sampling points were separated by a minimum distance of three meters. These samples were subsequently employed to analyze population structure, genetic diversity, gene flow, and admixture among them and within cultivars of the species.

**Figure 1.**
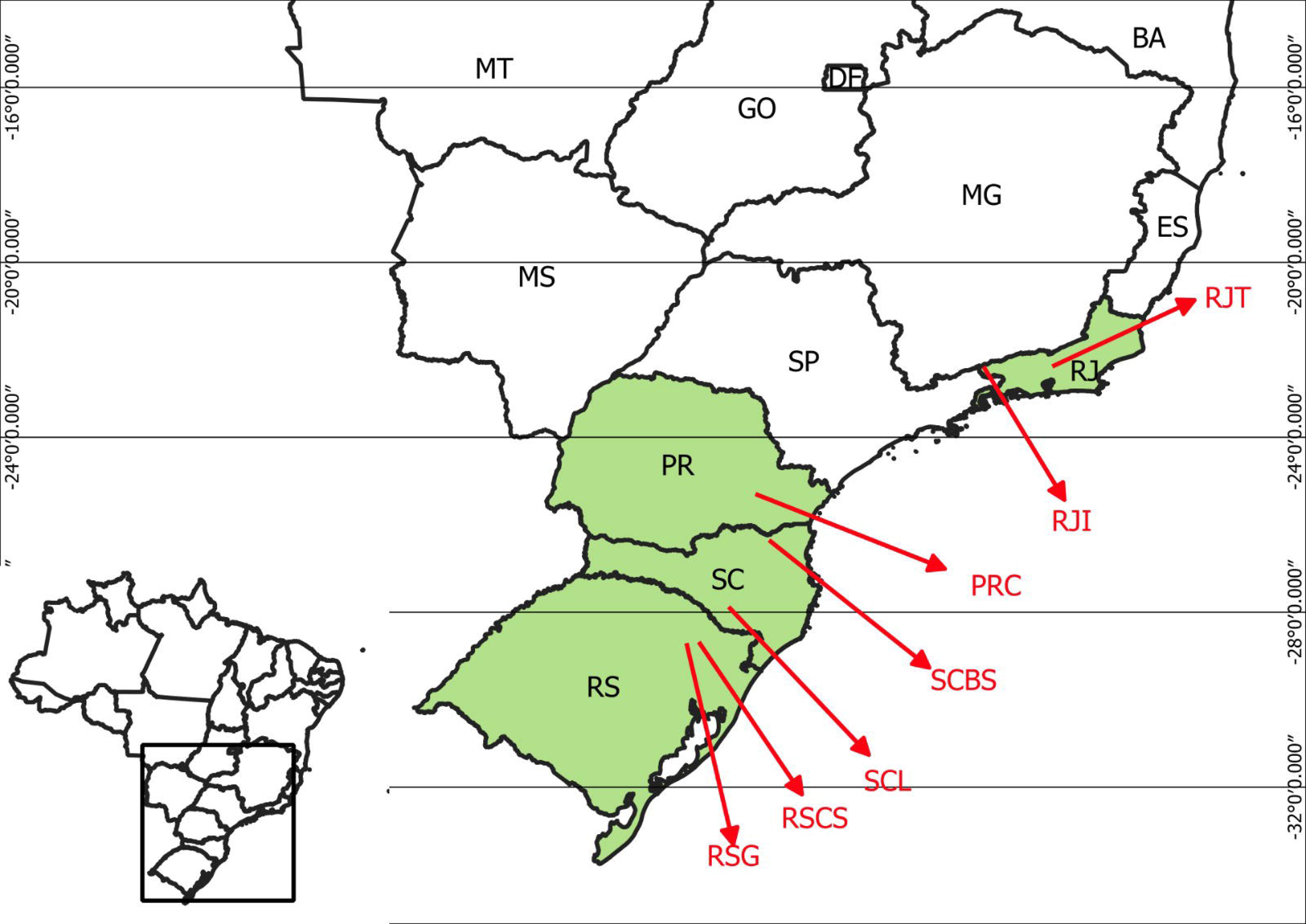
Map of the sample locations of naturalized populations of white clover in Brazil.

### Cultivars

Seeds of the cultivars BRS CGI/CGII, Ladino, Zapicán, Bombus, Jura, Merlyn, and Entrevero were procured commercially in 2021 (Table 1). Following germination and complete leaf expansion, fully developed leaves were carefully collected, individually stored in cloth sachets labeled with unique sample codes, and dried in silica gel until utilized for molecular analysis. Two samples per variety were used.

**Table 1.**
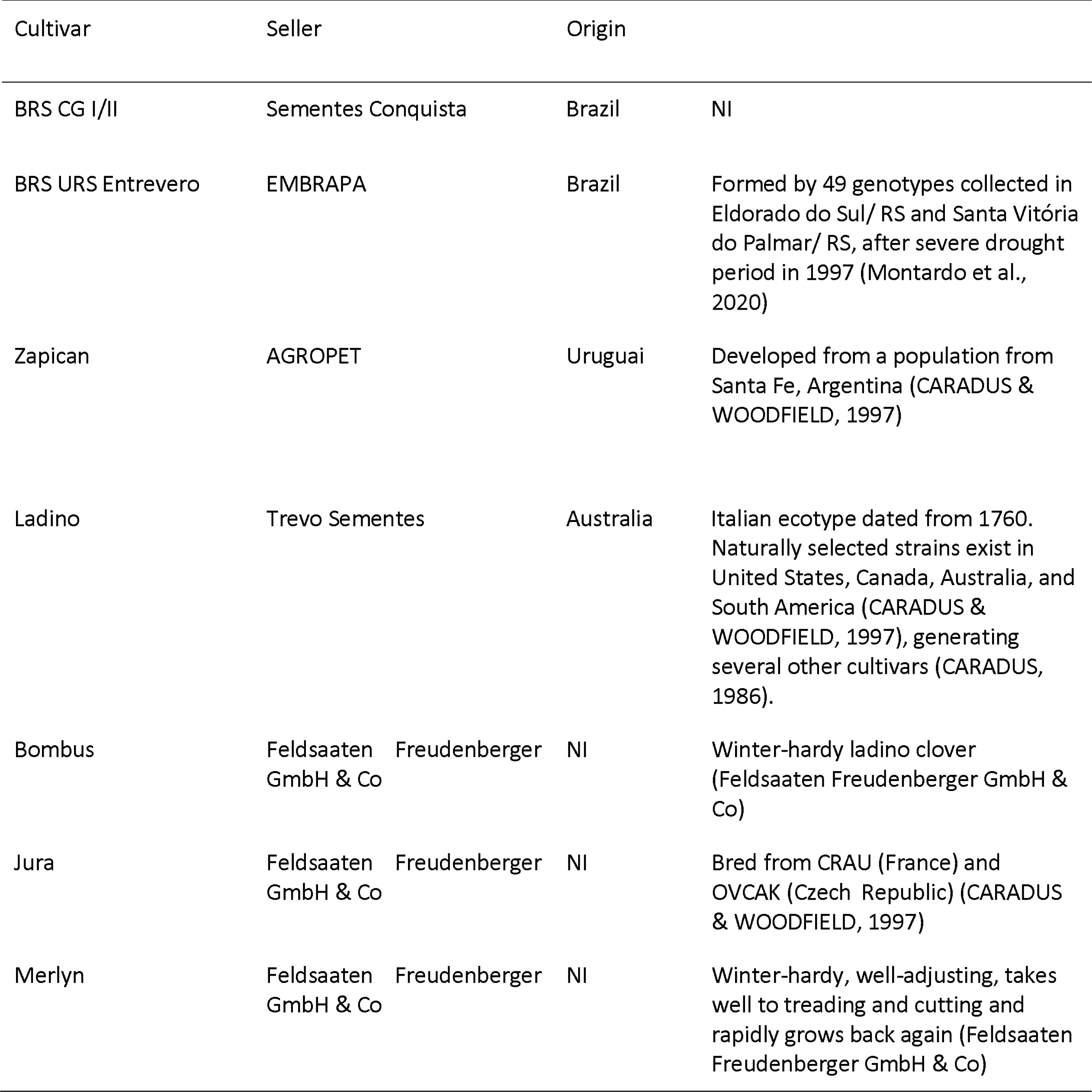
Cultivars of white clover obtained for molecular analyzes of admixture with Brazilian naturalized populations. NI: origin not informed by the manufacturer.

### DNA extraction and quantification

DNA extraction followed a modified protocol, combining the methods described originally by Agrawal et al. (2016) and Lira-Medeiros et al. (2015). A single leaf was crushed using metallic spheres, and then 500 μl of CTAB buffer (50 ml Tris 1M pH 8.0, 20 ml EDTA 0.5M pH 8.0, 41.5 g NaCl, 0.22 g ascorbic acid, 10 g CTAB with 500ml ddH2O) and 1 μl of β-mercaptoethanol were added. Samples were incubated in a water bath at 60°C for 20 minutes and centrifuged at 14,000 rpm for 3 minutes. The supernatant was transferred to a new tube, and an equal volume of chloroform was added, followed by centrifugation at maximum speed for 20 minutes. After moving the supernatant again, CIA (24:1 chloroform: isoamyl alcohol) was added and homogenized. Following another centrifugation and transfer of 150 μl of the supernatant to new tubes, 150 μl of ice-cold isopropanol was added, and the samples were incubated at −20°C for 20 minutes. The DNA was then precipitated by centrifugation, discarding the supernatant. The pellet was resuspended in 50 μl of deionized water and 3 μl of RNAse, incubated overnight at 37°C. Subsequently, 0.1 volume of 5M NaCl and two volumes of 100% ethanol were added. The samples were incubated in a freezer for 30 minutes, then centrifuged for 15 minutes at 12,000 rpm. The liquid phase was discarded, and 500 μl of 70% ethanol was added. The alcohol was discarded, and the pellet was resuspended in 50 μl of TE pH 8.0 buffer (10 mM Tris, 1 mM EDTA, and H2Od). A subset of samples was stored in the JBRJ DNA bank at −80°C as an ex-situ conservation measure. The samples were quantified using a Nanodrop spectrophotometer (ThermoFisher), and DNA integrity was verified by electrophoresis on a 1% agarose gel.

### MIG-seq protocol

The MIG-seq was performed according to the protocol in Suyama et al. (2022). Briefly, the MIG-seq library employed primer set-1 and the Suyama and Matsuki (2015) protocol, with specific modifications. These modifications included adjusting the temperature in the first PCR, incorporating purification/equalization steps, and eliminating short fragments (<250 bp) using AMPure XP. Sequencing was performed on an Illumina MiSeq platform using MiSeq Reagent Kit v3 (150 cycles) for 13 and 542 samples from a separate project. Paired-end and index sequencing covered both fragment and index ends. Notably, the “DarkCycle” option was modified to exclude the initial 17 bases (SSR and anchor regions) in both reads, and the effective read lengths were set at 80 bases for both reads in the software’s sample sheet.

### SNPs analysis

After merging the forward and reverse reads of each sample following Suyama and Matsuki (2015), low-quality reads were eliminated by using Trimmomatic v.0.32 (Bolger et al., 2014) with the following commands: HEADCROP:6, CROP:77, SLIDINGWINDOW:10:30, and MINLEN:51. Obtained reads were assembled using Stacks v.2.41 pipeline (Rochette et al. 2019). The minimum depth was set to 6 (-m 6), and default values were employed for the other option. Using populations program in Stacks, we extracted SNPs with genotyping rate > 80%. To remove paralogous loci and possible PCR errors, loci showing observed heterozygosity > 0.6 and SNPs with minor allele count < 0.3 were eliminated. We also filtered linked SNPs showing R^2^ > 0.4 using PLINK v1.90 (Chang et al. 2015).

The SNP data was analyzed using GeneAlex software v.6.5 (Peakall & Smouse 2006) to obtain the following diversity parameters: Shannon Information Index (I), expected and observed heterozygosity (He and Ho), number of alleles (Na), number of effective alleles (Ne), and percentage of polymorphic loci (P). Additionally, to assess genetic variation and differentiation among samples, F statistics and AMOVA were performed using the same software, excluding the “within individuals” option, and employing 9999 permutations for AMOVA. Also, the packages adegenet v.2.1.10 (Jombart and Ahmed 2011.), ade4 v.1.7-22 (Dray and Dufour 2007) and factoextra v.1.0.7 (Kassambara and Mundt 2020) were used in R v.4.2.3 (R Core Team 2023) for principal component analysis (PCA) and graphics. Population structure analysis was conducted using the ADMIXTURE v.1.3 software (Alexander et al. 2009). The number of ancestral populations (K) allowed varies from one to 10, with 10 replicate runs for each *K* value.

## Results

A total of 1097 SNPs were obtained from the analysis of seven Brazilian populations and seven cultivars. The results will be presented segmented, examining these groups separately and together.

### Brazilian naturalized populations

Initially, we tested the hypothesis of genetic structure of the populations in the southern (PR, SC, RS) and southeastern (RJ) regions, considering the substantial geographic distance between them (Table 2 - above diagonal) and the species reproductive characteristics in terms of pollination and limited dispersal. An AMOVA was performed with two regions to simulate this prediction. However, the results did not support our initial hypothesis, indicating 100% of the variation observed within populations for the species. Consequently, we conducted a new analysis, without regional cluster, resulting in 91% within-population variation (PhiPT = 0.095; p < 0.0001), indicating very low genetic structure.

**Table 2.**
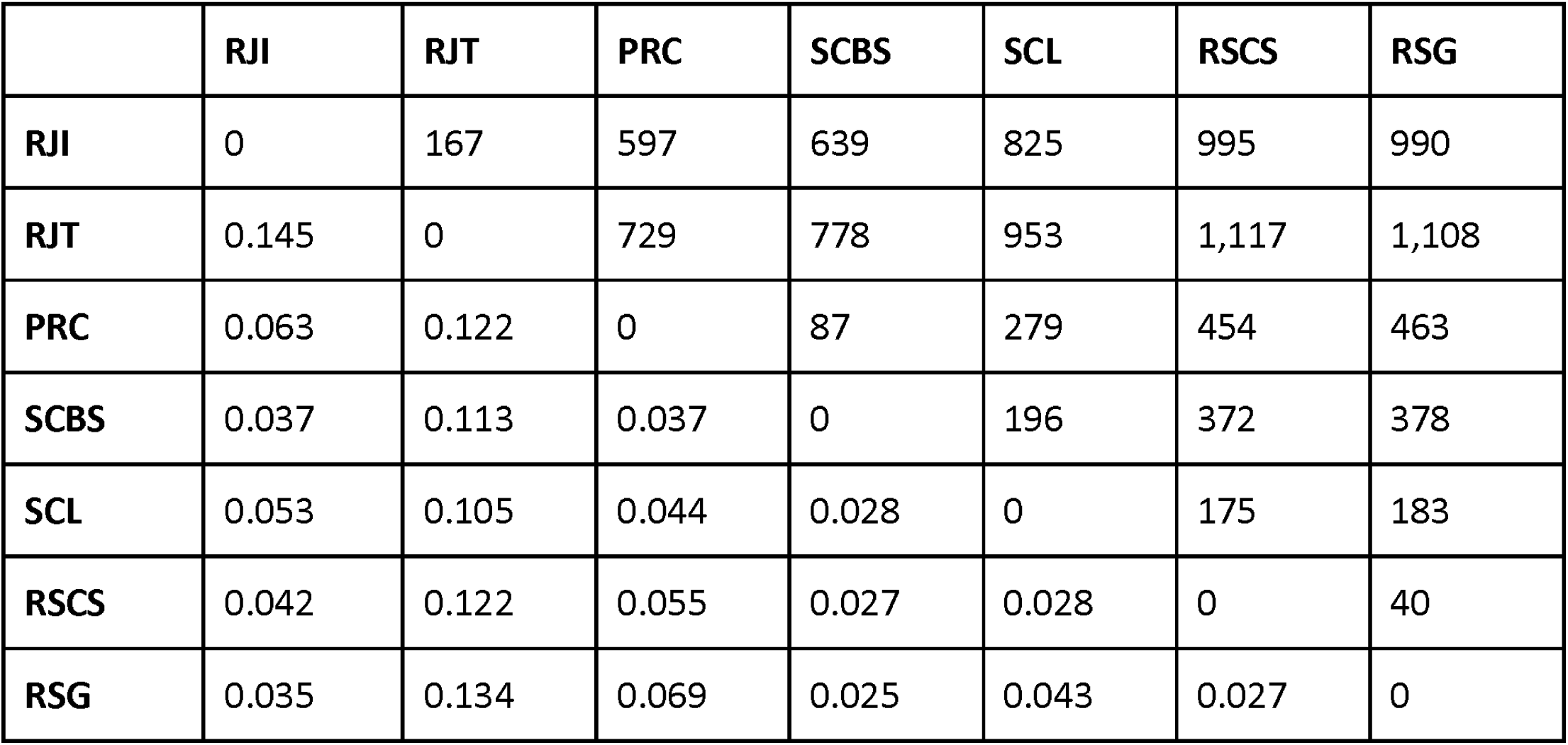
Pairwise Population Fst Values (lower diagonal) of Brazilian naturalized populations of white clover in Itatiaia (RJI), Teresópolis (RJT), Curitiba (PRC), São Bento do Sul (SCBS), Lages (SCL), Caxias do Sul (RSCS), and Gramado (RSG). Upper diagonal refers to approximatedly geographical distance (km) between municipality centroids.

Similarly, F statistics showed Fst = 0.062 (p < 0.0001). However, pairwise Fst, despite exhibiting low values, indicated moderate differentiation between two mountain cities geographically close to each other (0.145; RJI x RJT, and 0.134; RJT x RSG). We may observe that RJT has the highest values of genetic differentiation from all other populations, independently of geographical distance between them, ranging from 0.105 to 0.145 (Table 2). The lowest differentiation was between SCBS and RSG populations (0.025). All other populations had pairwaise Fst ranging from 0.025 to 0.069 (Table 2). The PCA shows that populations mostly overlapped, although several individuals from RJI, RJT, RSCS, RSG and PRC were dispersed (Figure 2). The populations PRC and RJT are likely more diverse since they had many individuals dispersed in PCA.

**Figure 2.**
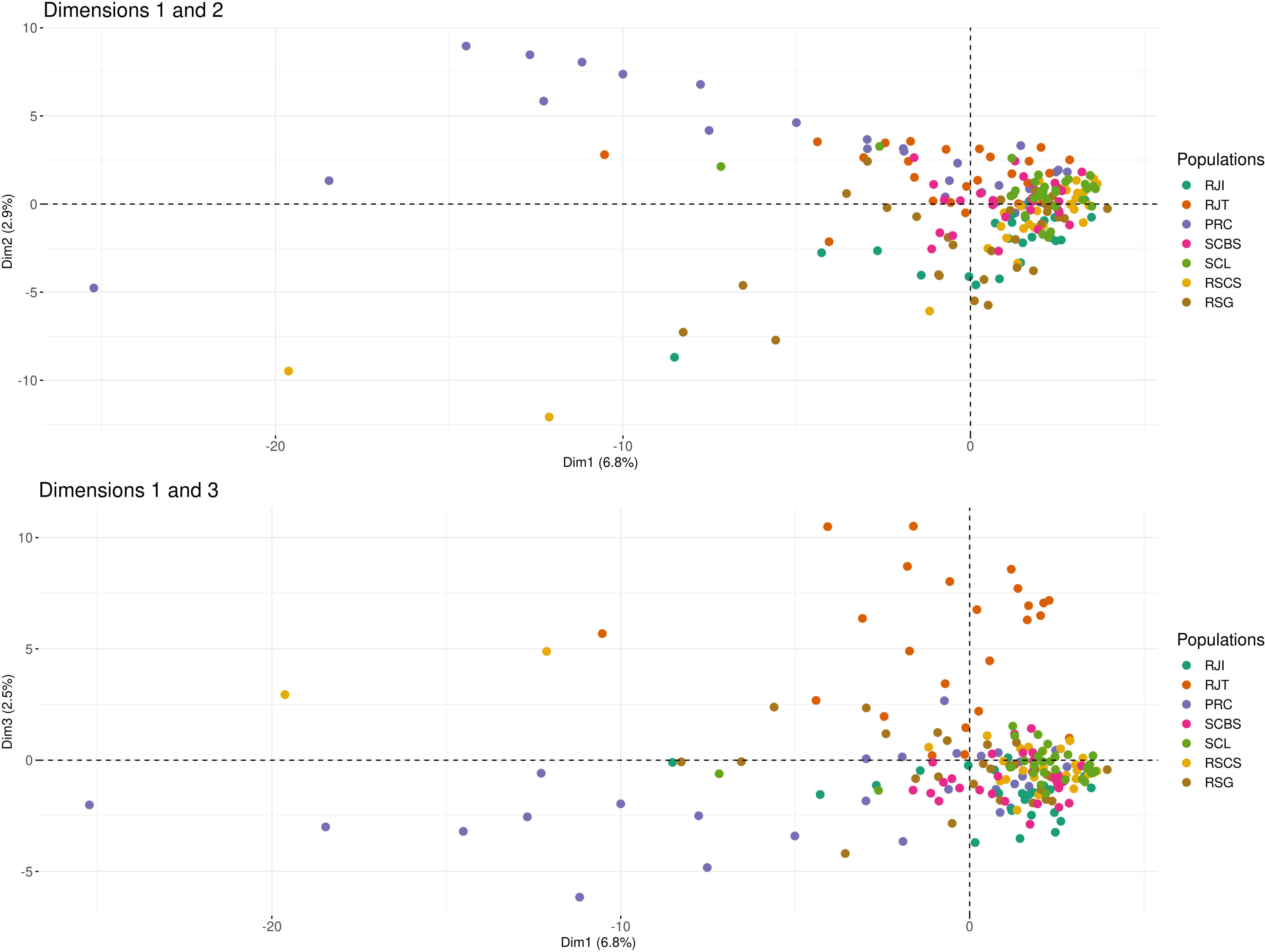
Principal Component Analysis of the genetic structure of Brazilian naturalized populations of white clover in the cities of Itatiaia (RJI), Teresópolis (RJT), Curitiba (PRC), São Bento do Sul (SCBS), Lages (SCL), Caxias do Sul (RSCS), and Gramado (RSG). Top figure showing dimensions 1 and 2, and bottom showing 1 and 3.

The overall genetic diversity statistics among naturalized populations revealed Na, Ne, and I as 1.578, 1.152, and 0.173, respectively. The Ho value (0.086) was lower than the He (0.102), with an inbreeding coefficient of 0.13. Percentage of polymorphic loci (P) was 57.82% (Table 3).

**Table 3.**
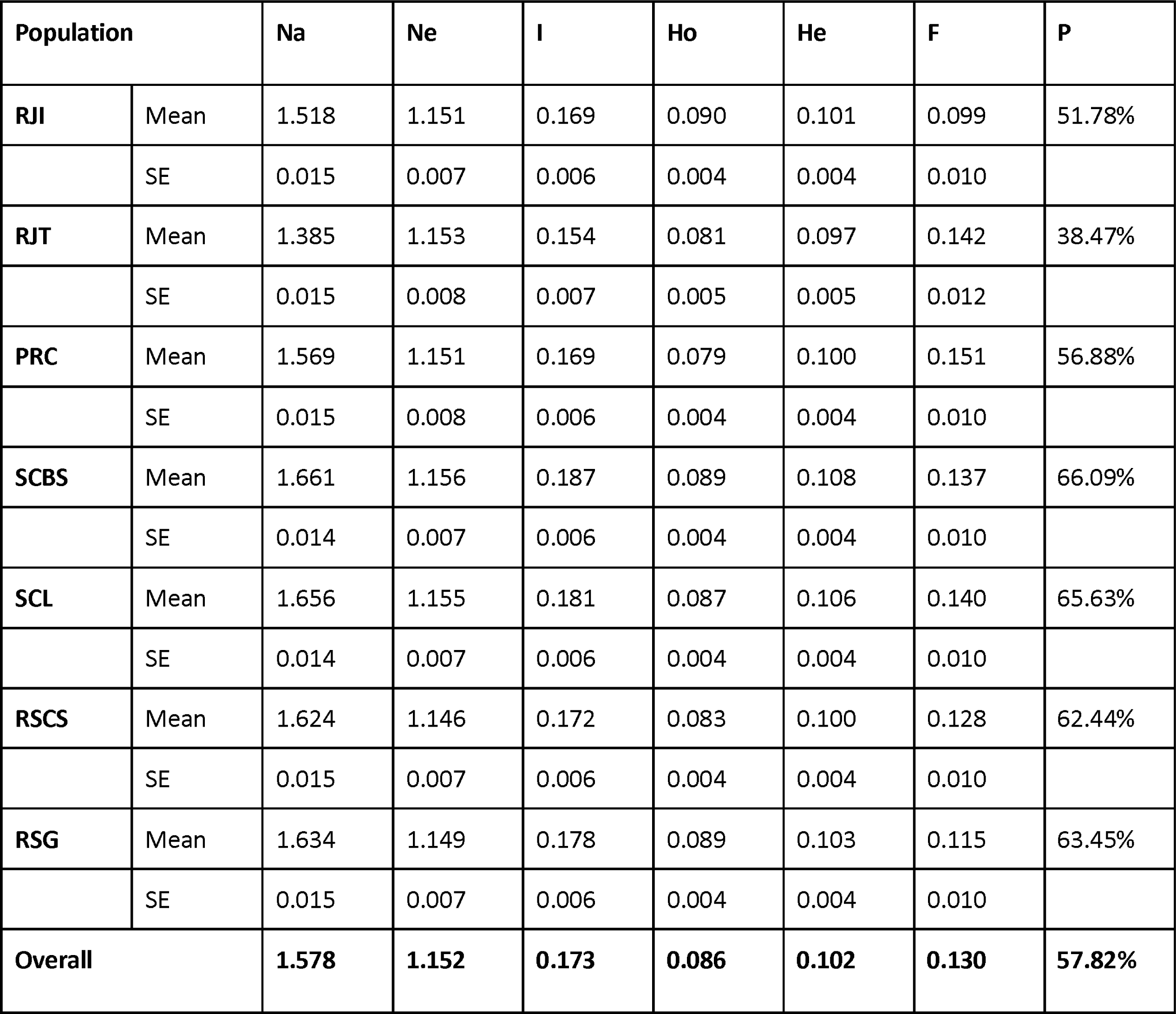
Genetic diversity indexes of the Brazilian naturalized populations of white clover in Itatiaia (RJI), Teresópolis (RJT), Curitiba (PRC), São Bento do Sul (SCBS), Lages (SCL), Caxias do Sul (RSCS), and Gramado (RSG). The indexes shown are: number of alleles (Na), effective number of alleles (Ne), Shannon diversity index (I), observed heterozygosity (Ho), expected heterozygosity (He), inbreeding coefficient (F) and percentage of polymorphic loci (P). SE: standard error.

| Population | | $N_a$ | $N_e$ | $I$ | $H_o$ | $H_e$ | $F$ | $P$ |
| --- | --- | --- | --- | --- | --- | --- | --- | --- |
| RJI | Mean | 1.518 | 1.151 | 0.169 | 0.090 | 0.101 | 0.099 | 51.78% |
|  | SE | 0.015 | 0.007 | 0.006 | 0.004 | 0.004 | 0.010 |  |
| RJT | Mean | 1.385 | 1.153 | 0.154 | 0.081 | 0.097 | 0.142 | 38.47% |
|  | SE | 0.015 | 0.008 | 0.007 | 0.005 | 0.005 | 0.012 |  |
| PRC | Mean | 1.569 | 1.151 | 0.169 | 0.079 | 0.100 | 0.151 | 56.88% |
|  | SE | 0.015 | 0.008 | 0.006 | 0.004 | 0.004 | 0.010 |  |
| SCBS | Mean | 1.661 | 1.156 | 0.187 | 0.089 | 0.108 | 0.137 | 66.09% |
|  | SE | 0.014 | 0.007 | 0.006 | 0.004 | 0.004 | 0.010 |  |
| SCL | Mean | 1.656 | 1.155 | 0.181 | 0.087 | 0.106 | 0.140 | 65.63% |
|  | SE | 0.014 | 0.007 | 0.006 | 0.004 | 0.004 | 0.010 |  |
| RSCS | Mean | 1.624 | 1.146 | 0.172 | 0.083 | 0.100 | 0.128 | 62.44% |
|  | SE | 0.015 | 0.007 | 0.006 | 0.004 | 0.004 | 0.010 |  |
| RSG | Mean | 1.634 | 1.149 | 0.178 | 0.089 | 0.103 | 0.115 | 63.45% |
|  | SE | 0.015 | 0.007 | 0.006 | 0.004 | 0.004 | 0.010 |  |
| Overall |  | 1.578 | 1.152 | 0.173 | 0.086 | 0.102 | 0.130 | 57.82% |

Analyzing the populations individually, Na ranged from 1.385 (RJT) to 1.661 (SCBS), Ne ranged from 1.146 (RSCS) to 1.156 (SCBS), I ranged from 0.154 (RJT) to 0.189 (SCBS). The largest difference between Ho and He and consequently higher inbreeding value was observed in PRC. The inbreeding coefficient (F) ranged from 0.099 (RJI) to 0.151 (PRC). The RJT population was the less diverse population reflected by the lowest P (38.47%; Table 3).

### Cultivars

In the PCA, international cultivars imported from Europe (Bombus, Jura, and Merlyn) are positioned in the right quadrant of the graphic, while the national and South American ones are in the left (Figure 3). The highest pairwise Fst was 0.313 between the Merlyn and Zapicán; the most similar were BRS CG I/II and Ladino (0.222) and BRS CG I/II and Entrevero (0.225) (Table 4). However, pairwise Fst values were high between cultivars.

**Figure 3.**
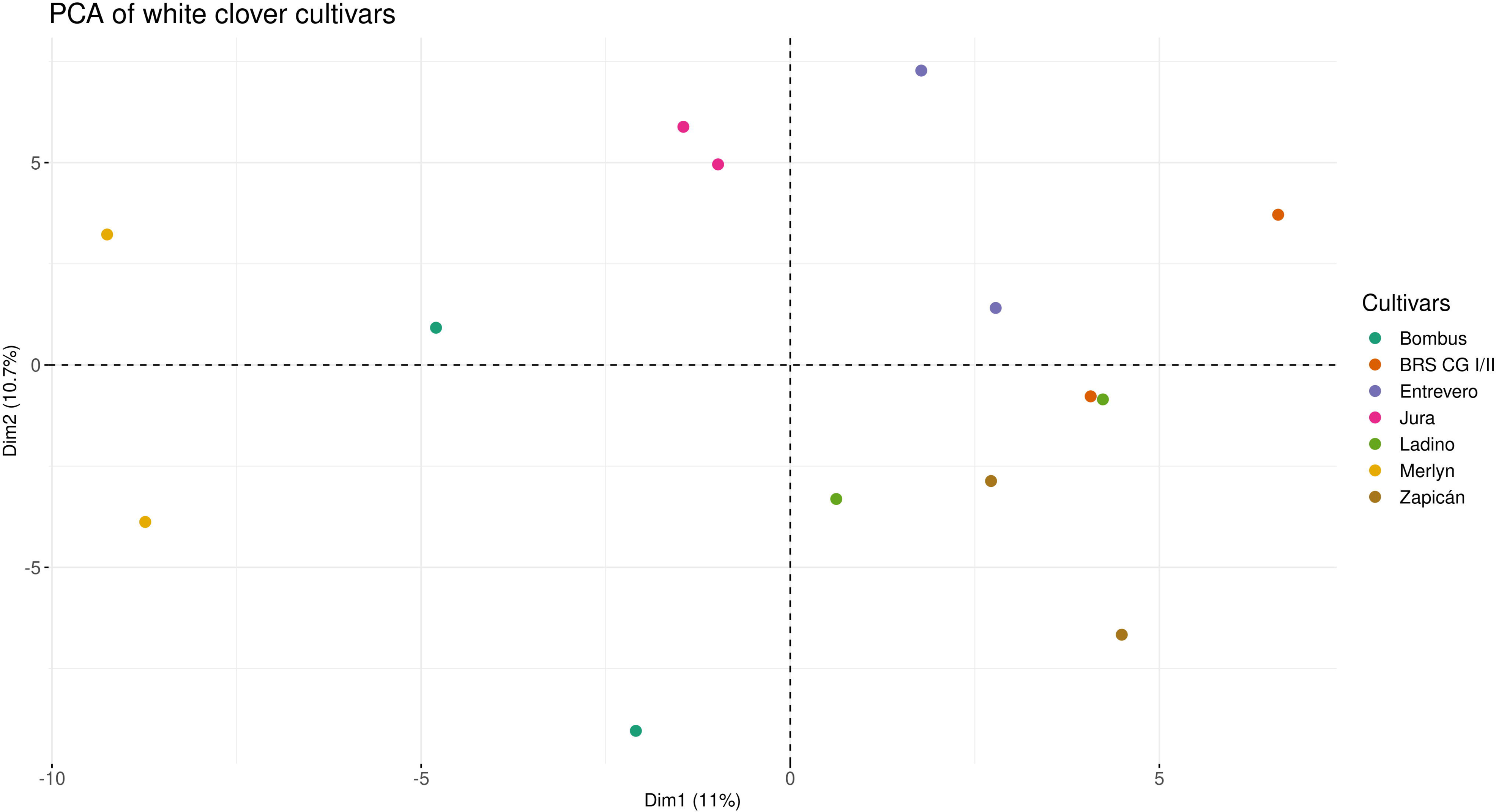
Principal Component Analysis of the genetic structure of cultivars of white clover: Bombus, BRS CG I/II, Entrevero, Jura, Ladino, Merlyn and Zapicán.

**Table 4.** Pairwise Population Fst Values of cultivars of white clover.

|  | Bombus | BRS CG I/II | Entrevero | Jura | Ladino | Merlyn | Zapican |
| --- | --- | --- | --- | --- | --- | --- | --- |
| Bombus | 0 |  |  |  |  |  |  |
| BRS CG I/II | 0.292 | 0 |  |  |  |  |  |
| Entrevero | 0.299 | 0.225 | 0 |  |  |  |  |
| Jura | 0.276 | 0.245 | 0.262 | 0 |  |  |  |
| Ladino | 0.268 | 0.222 | 0.258 | 0.262 | 0 |  |  |
| Merlyn | 0.259 | 0.292 | 0.276 | 0.258 | 0.272 | 0 |  |
| Zapican | 0.272 | 0.257 | 0.278 | 0.289 | 0.255 | 0.313 | 0 |

For the studied cultivars, Na was 1.138, Ne was 1.097, and I was 0.093. The Ho value (0.088) was higher than the He (0.064), resulting in a negative inbreeding coefficient of −0.363. Percentage of polymorphic loci (P) was very low (15.37%; Table 5), indicating high genetic homogeneity in cultivars, as expected.

**Table 5.**
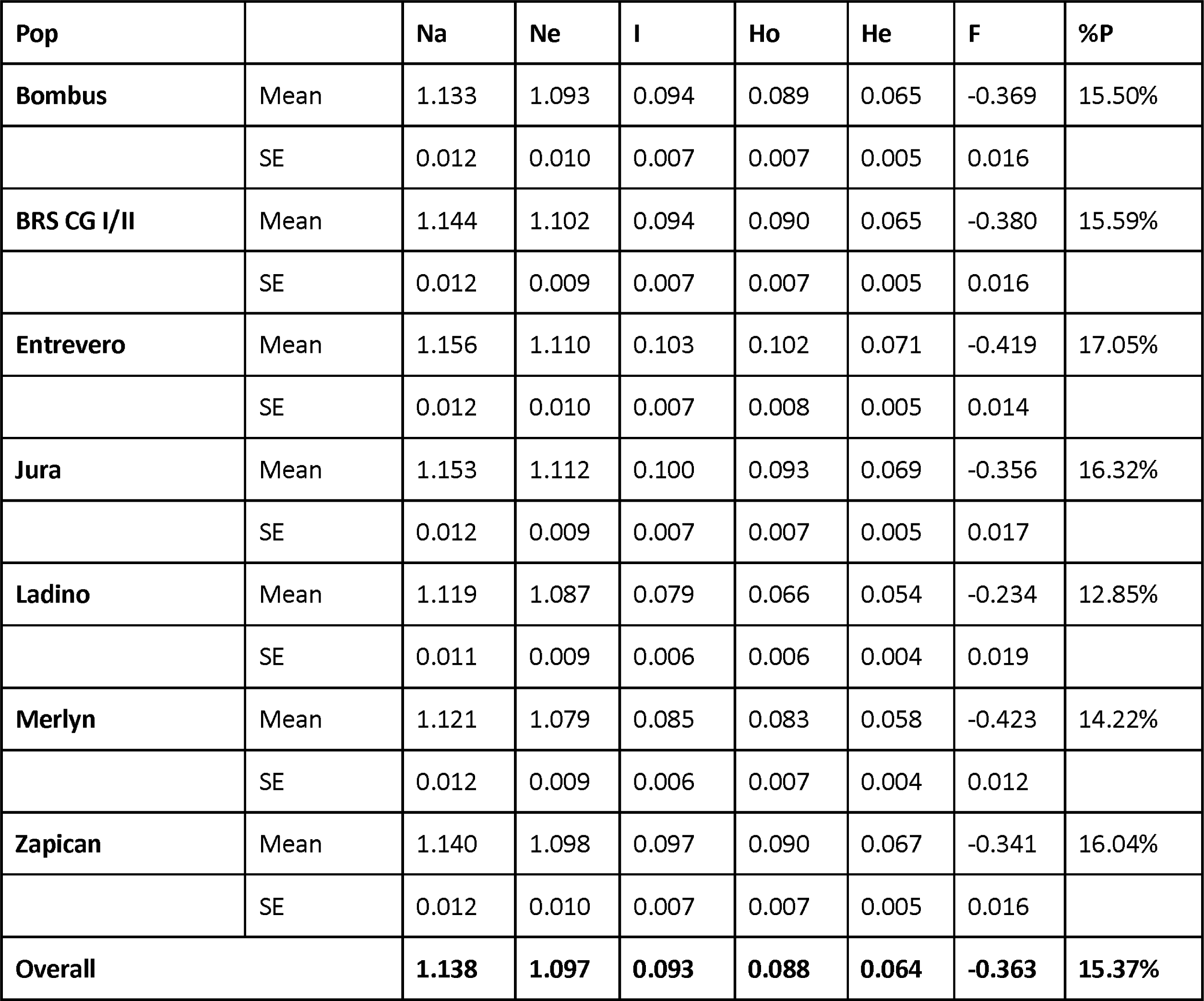
Genetic diversity indexes of cultivars of white clover. The indexes are number of alleles (Na), effective number of alleles (Ne), Shannon diversity index (I), observed heterozygosity (Ho), expected heterozygosity (He), inbreeding coefficient (F) and percentage of polymorphic loci (P). SE: standard error.

Analyzing the cultivars individually, Na ranged from 1.119 (Ladino) to 1.153 (Jura), Ne ranged from 1.079 (Merlyn) to 1.112 (Jura), and I ranged from 0.079 (Ladino) to 0.103 (Entrevero). Interestingly, Ho remained greater than He for all cultivars, and F was similarly negative for all cultivars, with the lowest value in Merlyn (−0.423). The Ladino variety had the lowest P as 12.85%, and Entrevro had the highest as 17.05% (Table 5).

### Overall genetic diversity of white clover

When analyzing the complete dataset, AMOVA revealed that 95% of the variation is within populations (PhiPT = 0.054; p < 0.0001), while F-statistics revealed 93% of variation within population (Fst = 0.071; p < 0.0001). The PCA clustered most samples, with some individuals from the PRC, RJI, RJT, RSCS, and RSG populations exhibiting higher genetic variability (Figure 4). Cultivars overlapped with most populations in PCA, possibly due to lower genetic variability of white clover in Brazil and some admixture between populations and cultivars. The admixture analysis indicated four clusters as optimal (Figure 5). The populations RJI and RSG clustered together with a unique genetic background, indicating similar ancestry. PRC individuals clustered in a second profile while RJT clustered in a third profile. Interestingly, cultivars showed mixed ancestry, as a result of a long improvement process. Additionally, SCBS showed mixed ancestry, and SCL and RSCS individuals had genetic profiles very similar to the cultivar profiles. The complete structure output for all tested K values is available in the Supplementary Material (Figure S2).

**Figure 4.**
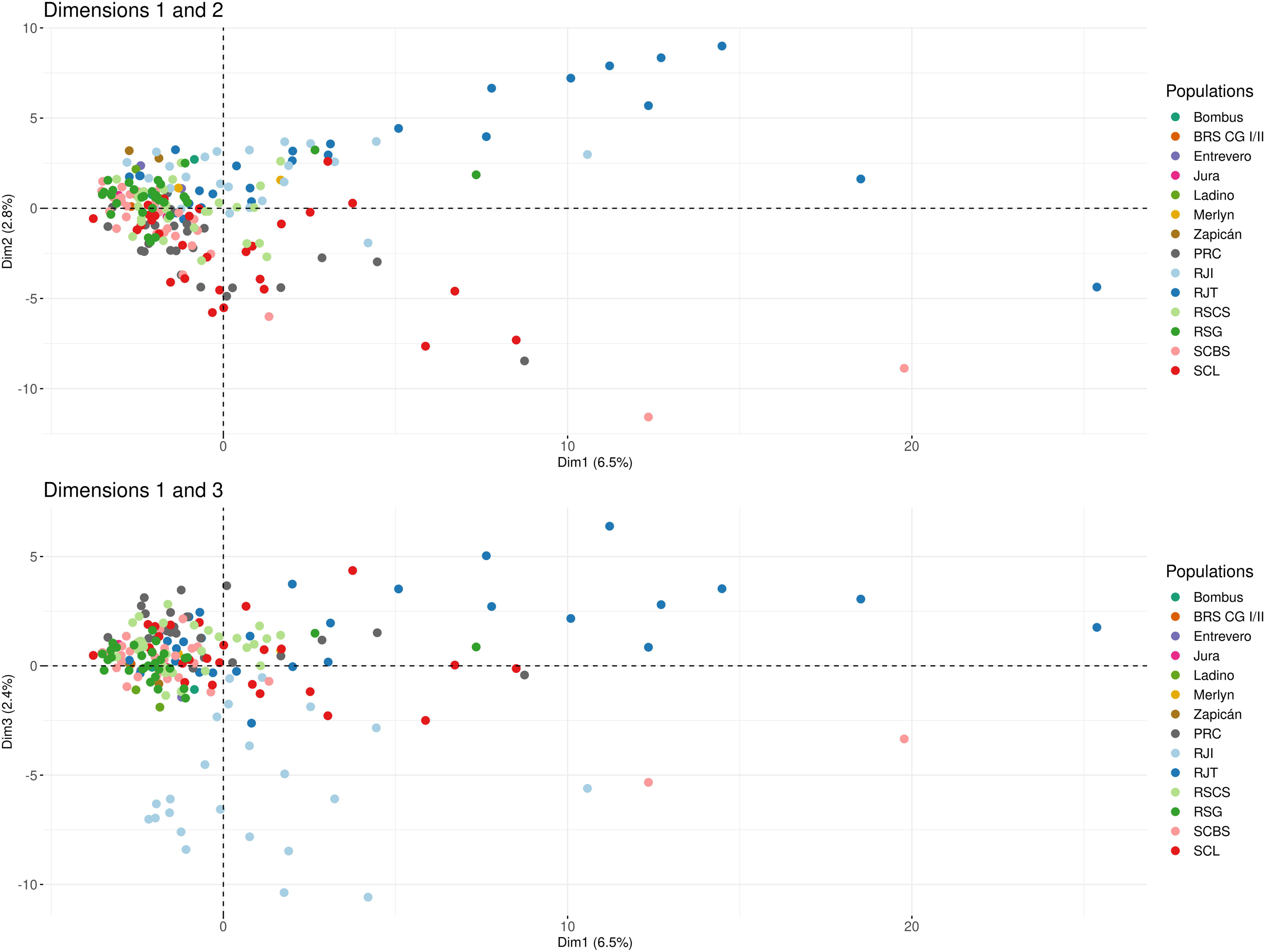
Principal component analysis (PCA) of the genetic structure of the cultivars Bombus, BRS CG I/II, Entrevero, Jura, Ladino, Merlyn and Zapicán, and Brazilian naturalized populations of white clover in the cities of Gramado (RSG), Caxias do Sul (RSCS), São Bento do Sul (SCBS), Lages (SCL), Curitiba (PRC), Teresópolis (RJT) and Itatiaia (RJI).

**Figure 5.**
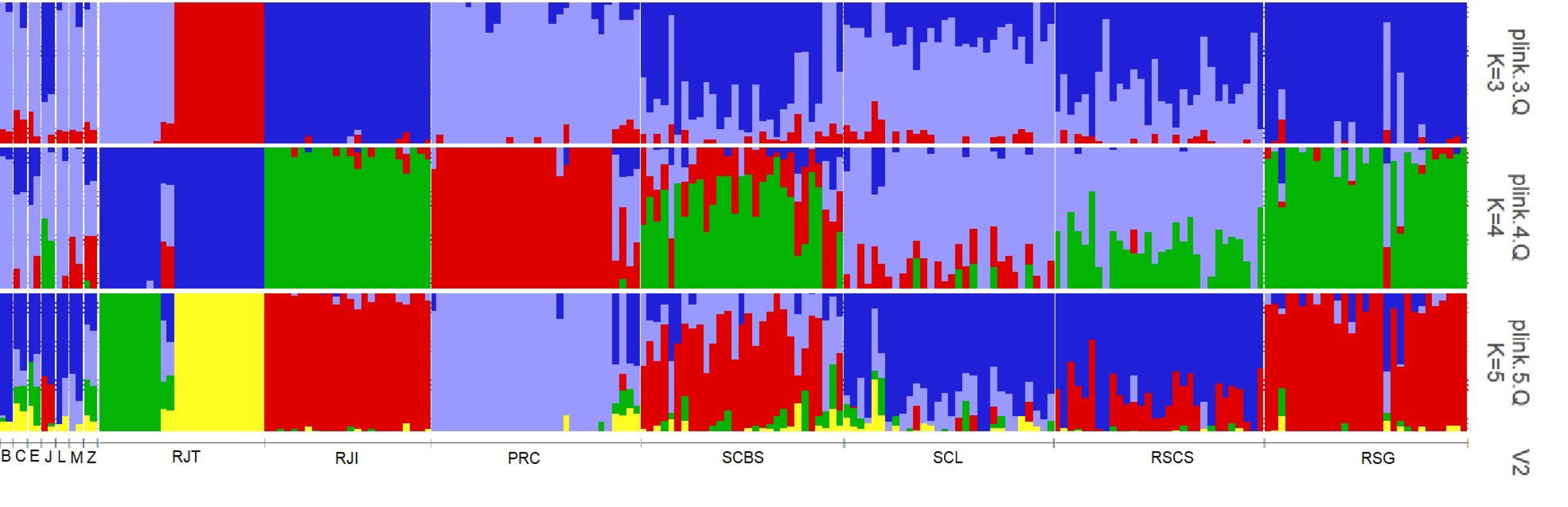
Genetic structure of the cultivars Bombus (B), BRS CG I/II (C), Entrevero (E), Jura (J), Ladino (L), Merlyn (M) and Zapicán (Z), and the Brazilian naturalized populations of white clover in Gramado (RSG), Caxias do Sul (RSCS), São Bento do Sul (SCBS), Lages (SCL), Curitiba (PRC), Teresópolis (RJT) and Itatiaia (RJI), with 3, 4 (optimal K) and 5 clusters.

## Discussion

### White clover in Brazil has low genetic diversity

The species’ colonization of new territories is a multifaceted process influenced by various factors, leaving a permanent mark on the genetic diversity of these newly established populations (Lachuth et al. 2010). Among the key determinants of success or failure in this process is the genetic diversity of the initial founding population, which is subsequently shaped by evolutionary forces and the species’ biological characteristics, such as reproductive and dispersal capabilities.

White clover, a species that exhibits mixed sexual and vegetative reproduction, is also self-incompatible. Meta-analyses indicate that self-incompatible species may be more susceptible to allelic richness loss, and herbaceous plants may exhibit lower genetic diversity due to factors such as life form (Gonzalez et al. 2020) and population size (Chung et al. 2020). However, studies have shown that outcrossing and short-lived perennial species typically would be more genetically diverse (Nybom and Bartish 2000; Watson-Jones et al. 2006; Hargreaves et al. 2010).

Global analyses of white clover accessions in introduced locations revealed an average Shannon diversity Index (*I)* of 0.427, similar to the values in the Americas (0.416) obtained with SSR markers (Wu et al. 2021). These results were higher than Korea’s and the United Kingdom’s natural populations in which *I* was 0.252 and 0.107, respectively (Hwang and Huh, 2016; Hargreaves et al. 2010). Interestingly, our dataset exhibited an average *I* of 0.173, lower than the previously reported for the Americas (germplasm) and Korean populations, and similar to the United Kingdom’s populations. A similar pattern was observed for P, with high average value in introduced locations (96.77%) and in the Americas accessions (95.31%; Wu et al. 2021). In contrast, our results yielded an average P of 57.82%, falling within the range between 42.1% observed in Korean populations (ISSR; Hwang and Huh, 2016) and 72% polymorphism in natural populations of northern Europe (Dabkevičienė et al. 2011). The previous *I* calculated for the Americas and non-native areas (Wu et al. 2021) was probably overestimated because they analyzed germplasm accessions without precise identification within the continent.

White clover cultivars showed similar genetic indexes among them and lower I and P compared to naturalized populations in Brazil. All tested cultivars displayed similar P rates, regardless of their origin. In contrast, other white clover cultivars reported in the literature have exhibited higher P using AFLP markers, ranging from 57 to 72% in the cultivars Norstar and Ramona (Collings et al. 2012) and an average of 89.5% in the cultivars silvit, jumbo, milligro, EmuNo.2, klondike, Huia, Haifa, Sulky, Purple and Ladino (Ma et al. 2020). Lower variability in this group was anticipated due to artificial selection and inbreeding during the improvement process, as confirmed by the negative F values obtained and low inbreeding coefficients. Therefore, although genetic diversity is low, the cultivars are divergent between themselves. Despite employing NGS methodology, the sample size may have influenced the lower polymorphism rates observed in our study. This suggests that analyzing a larger number of samples and accessions would be beneficial for cultivars genetic analysis, where greater homogeneity is expected due to artificial trait selection.

### Genetic structure of white clover in Brazil: ancestry and anthropogenic effect

Our findings indicate that most of the variability of white clover populations in Brazil is found within groups, in naturalized populations and cultivars. Global studies also indicated that 97% of the variability was present within populations, in germplasm banks from native and introduced localities (Wu et al. 2021). This concentration of intra-population variation is widely observed in local-scale studies, encompassing natural populations (Dabkevičienė et al. 2011; Collings et al, 2012; Hargreaves et al, 2010) and naturalized populations (Zhang et al. 2010; Pearson 2021; Hwang and Hu 2016; Kooyers and Olsen 2012), as well as cultivars (Ma et al. 2020; Nie et al. 2019; Khanlou et al. 2011).

In natural populations, genetic structure is strongly influenced by factors such as reproduction and dispersal patterns. As white clover reproduces sexually, the observation that nearly all genetic variability was found within populations and that there was no differentiation between populations suggests that gene flow is occurring without barriers for most populations.

In general, the genetic similarity of Brazilian populations could be attributed to a founder effect during their establishment. Also, herbaceous plants tend to have lower genetic diversity when compared to long living and arboreous plants. While there is no scientific literature explaining how white clover was brought in the country, it is possible to have occurred during the influx of European immigration to the southern region of Brazil in the 19th century or before. With a reduced initial genetic pool and low diversity in its original location the variability of the Brazilian populations would remain limited over time, resulting in high similarity between populations. In the ancestrality analysis, we can get some insights on the genetic pools that contributed to the formation of the populations analyzed. RSCS has a similar combination of genotypes to the European cultivar Jura, and SCL with cultivars from South America, indicating that the individuals from these cities may have originated from different cultivars. The SCBS seems to be the most ancestry-mixed population, which might indicate that different European native pools contributed to its formation. Also, we can observe some individuals in SCL which are similar to SCBS, possibly due to the proximity of these two cities.

Controversially, we can observe some cases in which the population’s origin is based on different genotypes. Those are the cases of PRC and RJT populations in comparison to the other locations. Interestingly, the RJT population was divided into three homogeneous subgroups (Figure 1). Combining the PCA information with sample location data, we confirmed that these subgroups were composed of: 1) samples collected randomly from various locations within the city that were similar to Latin American cultivars’ genotypes, 2) samples from an agricultural area, and 3) samples from a private property. So, the divergent clustering of RJT individuals is probably due to unintentional anthropogenic effects.

PRC individuals had other ancestry, not similar to any other location, although some PRC samples have similar genetic background as of analyzed cultivars. Cases of population differentiation have only been observed when associated with geographic isolation, as in the United Kingdom. There, more distant islands became strongholds of species diversity, while on the mainland, a tendency towards homogenization and similarity between populations was more likely (Hargreaves et al. 2010). The authors further point out that in the absence of long-distance dispersal in this species over sea barriers, anthropogenic dispersal will have contributed to the genetic similarity observed. The cases of RJT and PRC are not necessarily a matter of geographic isolation, but might be way more related to the origin of the founding populations, that might have come from one specific native location for each site or one specific cultivar (not analyzed here).

On the other hand, RJI and RSG had the same ancestry and may reflect a European introduction not related to its usage as forage. Possibly, white clover was introduced for landscaping while Europeans were in the mountainous regions in Brazil, where they greatly influenced some cities’ culture. These two populations and part of SCBC population are likely to have similar origin due to ancestry genotypes, and no relationship with cultivar’s genotypic diversity.

For the cultivars analyzed, some divergence was observed between national/South American cultivars and those sold in Europe. This can be attributed to the selection of strains specifically tailored to meet the environmental demands of each region. However, depending on the cultivars analyzed, no variation may be observed between them especially due to similar parental origin of several accessions (Ma et al. 2020).

The lack of differentiation is also evident when comparing cultivars and naturalized populations. This data suggests that even with improvement for cultivation, there has not been sufficient time for these groups to differentiate genetically in Brazil. This finding highlights the potential of cultivars to colonize the territory in cases of agricultural escape and cause a reduction of genetic variability in naturalized populations due to homogenization.

### 3) White clover: is it showing its colors?

For a plant to successfully undergo an invasion process, it must navigate through distinct phases: introduction, naturalization and invasion (Richardson et al. 2000). The ability to progress through these phases is contingent upon both ecological factors, such as dispersal capacity, reproductive mode, niche compatibility, and genetic factors. It is well established in the literature that greater genetic diversity enhances colonization success (Crawford and Whitney 2010, Sakai et al. 2001).

While white clover is considered invasive in some countries (Meneer et al. 2005; Golding 2003; MDC 2023; Guy et al. 2013), in Brazil it is considered naturalized (*Trifolium repens*; Iganci et al., 2024). In field observations, white clover is only found in grassy areas or roadsides, restricted to regions with milder or mountainous climates. Moreover, due to the substantial portion of Brazil’s territory lying within the tropical belt, the species’ range of occurrence is climatically limited.

Brazilian populations exhibit lower genetic diversity than in other introduced locations, but similar to UK diversity levels. The literature does not conclude about the level of genetic diversity presented by invasive species, if they are greater in general (Lachmuth et al. 2010) or if invasive clonal species are a particular case and present lower levels (González et al. 2016). Considering the significant contribution of sexual reproduction for clover’s population structure, its introduction during European immigration over 100 years ago, and still presenting low genetic diversity, white clover does not align with the genetic and ecological characteristics of an invasive species in Brazil. Therefore, its current classification as a naturalized species is accurate since white clover’s behavior and genetic profile do not align with those of an invasive species.

## Conclusions

This is the first Brazilian study regarding white clover origins within the national territory and that was able to confirm that its classification as naturalized is appropriate. The results presented here can be used as a startpoint to foment other studies on its origins and dispersion.

Despite the similarity of white clover populations across the country and their levels of genetic variability, they have different ancestries. It is possible that part of the populations were originally formed with genetic pool from native areas, brought during European immigration such as in mountainous locations. However, parts of the populations have the same cultivar ancestry, indicating that cultivars might be escaping to naturalized areas instead of staying limited to forage areas. As white clover has biotechnological applications as well, then monitoring its behavioral and genetic profile over time is crucial to prevent stronger admixture between them, causing harm to both naturalized populations (due to homogenization) and to cultivars (due to loss of interesting traits).

In a broader look, it was possible to highlight the importance of genetic studies in classifying species, a point that does not receive proper attention in many projects. With this type of data, it is possible to identify characteristic events that could potentially harm biodiversity, which is not always feasible using other types of studies, such as ecological data.

## Acknowledgments

The Conselho Nacional de Desenvolvimento Científico e Tecnológico (CNPq) has supported this work. We also thank Nathalia Brandão and Vitor G. Cunha for the map’s graphic production.

