## Supplementary Table S1 for "Insights from Genetic Studies: SNP Analyses Confirm White Clover Naturalization in Brazil"

Table S1: Brazilian naturalized white clover samples collected in Gramado (RSG), Caxias do Sul (RSCS), São Bento do Sul (SCBS), Lages (SCL), Curitiba (PRC), Teresópolis (RJT) and Itatiaia (RJI).

| POP ID | SAMPLE ID | LATITUDE | LONGITUDE | ELEVATION |
|---|---|---|---|---|
| Gramado | RSG101 | -29.365161 | -50.803550 | 830 |
| Gramado | RSG102 | -29.364050 | -50.802366 | 833 |
| Gramado | RSG103 | -29.364015 | -50.803428 | 838 |
| Gramado | RSG104 | -29.365296 | -50.803465 | 831 |
| Gramado | RSG109 | -29.364471 | -50.803770 | 828 |
| Gramado | RSG110 | -29.365082 | -50.803526 | 828 |
| Gramado | RSG111 | -29.364049 | -50.809278 | 851 |
| Gramado | RSG112 | -29.363634 | -50.808941 | 842 |
| Gramado | RSG113 | -29.363926 | -50.809562 | 842 |
| Gramado | RSG114 | -29.363661 | -50.809854 | 839 |
| Gramado | RSG115 | -29.363996 | -50.809362 | 841 |
| Gramado | RSG117 | -29.363948 | -50.809106 | 846 |
| Gramado | RSG118 | -29.363476 | -50.809075 | 833 |
| Gramado | RSG119 | -29.363643 | -50.809093 | 834 |
| Gramado | RSG120 | -29.363673 | -50.809516 | 842 |
| Gramado | RSG122 | -29.359970 | -50.827338 | 828 |
| Gramado | RSG126 | -29.360606 | -50.848399 | 825 |
| Gramado | RSG128 | -29.360311 | -50.848540 | 825 |
| Gramado | RSG130 | -29.359826 | -50.855273 | 820 |
| Gramado | RSG132 | -29.386656 | -50.884796 | 823 |
| Gramado | RSG134 | -29.393899 | -50.877165 | 841 |
| Gramado | RSG136 | -29.359519 | -50.855990 | 823 |
| Gramado | RSG137 | -29.359211 | -50.854832 | 821 |
| Gramado | RSG139 | -29.359583 | -50.855232 | 819 |
| Gramado | RSG141 | -29.394638 | -50.877585 | 855 |
| Gramado | RSG146 | -29.386710 | -50.883819 | 833 |
| Gramado | RSG147 | -29.394562 | -50.877887 | 848 |
| Gramado | RSG148 | -29.393130 | -50.875820 | 853 |
| Gramado | RSG150 | -29.385060 | -50.877135 | 816 |
| Caxias do Sul | RSCS155 | -29.172833 | -51.179097 | 753 |
| Caxias do Sul | RSCS160 | -29.172979 | -51.179243 | 745 |
| Caxias do Sul | RSCS161 | -29.172410 | -51.179119 | 748 |
| Caxias do Sul | RSCS166 | -29.172113 | -51.178390 | 753 |
| Caxias do Sul | RSCS167 | -29.172575 | -51.178846 | 743 |
| Caxias do Sul | RSCS168 | -29.172329 | -51.179000 | 743 |
| Caxias do Sul | RSCS171 | -29.160096 | -51.156401 | 815 |
| Caxias do Sul | RSCS172 | -29.160375 | -51.156500 | 823 |
| Caxias do Sul | RSCS175 | -29.170841 | -51.191923 | 783 |
| Caxias do Sul | RSCS178 | -29.170015 | -51.196086 | 774 |
| Caxias do Sul | RSCS179 | -29.169967 | -51.196277 | 770 |
| Caxias do Sul | RSCS180 | -29.169770 | -51.196193 | 770 |
| Caxias do Sul | RSCS181 | -29.169681 | -51.195764 | 775 |
| Caxias do Sul | RSCS183 | -29.162040 | -51.188877 | 743 |
| Caxias do Sul | RSCS184 | -29.162008 | -51.190179 | 740 |
| Caxias do Sul | RSCS185 | -29.161709 | -51.190481 | 746 |
| Caxias do Sul | RSCS186 | -29.163484 | -51.188556 | 765 |
| Caxias do Sul | RSCS187 | -29.163696 | -51.188504 | 755 |
| Caxias do Sul | RSCS188 | -29.161662 | -51.189897 | 734 |
| Caxias do Sul | RSCS189 | -29.161659 | -51.189104 | 730 |
| Caxias do Sul | RSCS190 | -29.161808 | -51.189237 | 728 |
| Caxias do Sul | RSCS191 | -29.162656 | -51.150662 | 824 |
| Caxias do Sul | RSCS192 | -29.163653 | -51.149970 | 822 |
| Caxias do Sul | RSCS193 | -29.162808 | -51.150235 | 823 |
| Caxias do Sul | RSCS194 | -29.163560 | -51.150333 | 824 |
| Caxias do Sul | RSCS195 | -29.162807 | -51.150447 | 823 |
| Caxias do Sul | RSCS196 | -29.163616 | -51.150485 | 825 |
| Caxias do Sul | RSCS197 | -29.163418 | -51.149795 | 820 |
| Caxias do Sul | RSCS198 | -29.162840 | -51.150356 | 823 |
| Caxias do Sul | RSCS199 | -29.162583 | -51.150851 | 826 |
| Lages | SCL201 | -27.819827 | -50.262600 | 918 |
| Lages | SCL202 | -27.819708 | -50.261801 | 915 |
| Lages | SCL206 | -27.819749 | -50.260769 | 920 |
| Lages | SCL207 | -27.819708 | -50.260382 | 921 |
| Lages | SCL208 | -27.816917 | -50.267083 | 939 |
| Lages | SCL209 | -27.816969 | -50.266705 | 915 |
| Lages | SCL210 | -27.815357 | -50.280124 | 912 |
| Lages | SCL213 | -27.814721 | -50.285236 | 925 |
| Lages | SCL215 | -27.814396 | -50.285525 | 924 |
| Lages | SCL216 | -27.813631 | -50.285420 | 922 |
| Lages | SCL217 | -27.813687 | -50.285739 | 920 |
| Lages | SCL219 | -27.812478 | -50.285288 | 911 |
| Lages | SCL220 | -27.812397 | -50.285448 | 907 |
| Lages | SCL222 | -27.806029 | -50.308179 | 905 |
| Lages | SCL223 | -27.810361 | -50.301696 | 907 |
| Lages | SCL225 | -27.805935 | -50.307593 | 907 |
| Lages | SCL226 | -27.806557 | -50.308090 | 904 |
| Lages | SCL232 | -27.811784 | -50.322153 | 900 |
| Lages | SCL233 | -27.813727 | -50.322986 | 899 |
| Lages | SCL234 | -27.812962 | -50.323140 | 913 |
| Lages | SCL236 | -27.810938 | -50.321237 | 901 |
| Lages | SCL239 | -27.813800 | -50.323915 | 905 |
| Lages | SCL240 | -27.810905 | -50.320764 | 886 |
| Lages | SCL243 | -27.819445 | -50.321336 | 884 |
| Lages | SCL244 | -27.814362 | -50.318642 | 889 |
| Lages | SCL245 | -27.814507 | -50.318592 | 882 |
| Lages | SCL247 | -27.815795 | -50.318370 | 884 |
| Lages | SCL248 | -27.814196 | -50.318887 | 887 |
| Lages | SCL249 | -27.819730 | -50.321510 | 887 |
| Lages | SCL250 | -27.816185 | -50.318782 | 883 |

| | | | | |
|---|---|---|---|---|
| São Bento do Sul | SCBS252 | -26.220208 | -49.345920 | 893 |
| São Bento do Sul | SCBS253 | -26.220294 | -49.346859 | 894 |
| São Bento do Sul | SCBS254 | -26.217479 | -49.347036 | 874 |
| São Bento do Sul | SCBS256 | -26.222826 | -49.346293 | 902 |
| São Bento do Sul | SCBS259 | -26.220453 | -49.346041 | 891 |
| São Bento do Sul | SCBS262 | -26.227259 | -49.347682 | 926 |
| São Bento do Sul | SCBS265 | -26.223836 | -49.346697 | 898 |
| São Bento do Sul | SCBS266 | -26.223727 | -49.347435 | 902 |
| São Bento do Sul | SCBS269 | -26.246823 | -49.351442 | 848 |
| São Bento do Sul | SCBS270 | -26.245692 | -49.351665 | 838 |
| São Bento do Sul | SCBS271 | -26.245276 | -49.354967 | 865 |
| São Bento do Sul | SCBS272 | -26.245343 | -49.354426 | 864 |
| São Bento do Sul | SCBS275 | -26.240776 | -49.396124 | 871 |
| São Bento do Sul | SCBS276 | -26.246113 | -49.355761 | 868 |
| São Bento do Sul | SCBS278 | -26.244016 | -49.354337 | 857 |
| São Bento do Sul | SCBS279 | -26.240686 | -49.396553 | 872 |
| São Bento do Sul | SCBS280 | -26.246045 | -49.356011 | 867 |
| São Bento do Sul | SCBS281 | -26.240847 | -49.397856 | 876 |
| São Bento do Sul | SCBS282 | -26.239538 | -49.398370 | 855 |
| São Bento do Sul | SCBS285 | -26.240610 | -49.397864 | 874 |
| São Bento do Sul | SCBS287 | -26.237148 | -49.402917 | 857 |
| São Bento do Sul | SCBS288 | -26.239564 | -49.398599 | 856 |
| São Bento do Sul | SCBS289 | -26.239489 | -49.397225 | 865 |
| São Bento do Sul | SCBS291 | -26.229325 | -49.410686 | 882 |
| São Bento do Sul | SCBS292 | -26.229101 | -49.411064 | 879 |
| São Bento do Sul | SCBS294 | -26.229466 | -49.413432 | 850 |
| São Bento do Sul | SCBS295 | -26.229449 | -49.413532 | 860 |
| São Bento do Sul | SCBS296 | -26.234875 | -49.401292 | 840 |
| São Bento do Sul | SCBS298 | -26.234004 | -49.401435 | 846 |
| São Bento do Sul | SCBS299 | -26.234616 | -49.401472 | 840 |
| Curitiba | PRC301 | -25.431543 | -49.310599 | 898 |
| Curitiba | PRC302 | -25.431389 | -49.310257 | 900 |
| Curitiba | PRC303 | -25.428276 | -49.305637 | 897 |
| Curitiba | PRC307 | -25.428685 | -49.306249 | 901 |
| Curitiba | PRC308 | -25.431894 | -49.311938 | 916 |
| Curitiba | PRC311 | -25.380304 | -49.280803 | 976 |
| Curitiba | PRC314 | -25.378967 | -49.282034 | 976 |
| Curitiba | PRC315 | -25.378953 | -49.282345 | 973 |
| Curitiba | PRC317 | -25.378616 | -49.280165 | 980 |
| Curitiba | PRC319 | -25.380305 | -49.281201 | 977 |
| Curitiba | PRC320 | -25.380605 | -49.280374 | 983 |
| Curitiba | PRC321 | -25.508300 | -49.205785 | 879 |
| Curitiba | PRC322 | -25.511199 | -49.206175 | 880 |
| Curitiba | PRC323 | -25.510912 | -49.206011 | 880 |
| Curitiba | PRC326 | -25.509117 | -49.205681 | 880 |
| Curitiba | PRC330 | -25.508333 | -49.205047 | 884 |
| Curitiba | PRC331 | -25.525445 | -49.223245 | 880 |
| Curitiba | PRC332 | -25.525285 | -49.223057 | 878 |
| Curitiba | PRC333 | -25.526348 | -49.222854 | 878 |
| Curitiba | PRC338 | -25.525233 | -49.223373 | 877 |
| Curitiba | PRC339 | -25.527473 | -49.220607 | 880 |
| Curitiba | PRC340 | -25.525663 | -49.223002 | 877 |
| Curitiba | PRC341 | -25.404467 | -49.303945 | 906 |
| Curitiba | PRC343 | -25.404302 | -49.304048 | 906 |
| Curitiba | PRC344 | -25.404344 | -49.303521 | 904 |
| Curitiba | PRC345 | -25.405376 | -49.304094 | 902 |
| Curitiba | PRC346 | -25.395375 | -49.305737 | 902 |
| Curitiba | PRC347 | -25.395234 | -49.305894 | 901 |
| Curitiba | PRC348 | -25.395532 | -49.306036 | 905 |
| Curitiba | PRC349 | -25.395326 | -49.305464 | 906 |
| Teresópolis | RJT3 | -22.44606 | -42.981889 | 932 |
| Teresópolis | RJT32 | -22.353312 | -42.869291 | 830 |
| Teresópolis | RJT33 | -22.306789 | -42.992899 | 1,064 |
| Teresópolis | RJT34 | -22.306791 | -42.992899 | 1,064 |
| Teresópolis | RJT35 | -22.306787 | -42.992907 | 1,064 |
| Teresópolis | RJT36 | -22.306756 | -42.992913 | 1,065 |
| Teresópolis | RJT37 | -22.306764 | -42.992918 | 1,065 |
| Teresópolis | RJT38 | -22.306764 | -42.992918 | 1,065 |
| Teresópolis | RJT39 | -22.306764 | -42.992918 | 1,066 |
| Teresópolis | RJT41 | -22.306776 | -42.992932 | 1,064 |
| Teresópolis | RJT42 | -22.306768 | -42.992930 | 1,065 |
| Teresópolis | RJT43 | -22.306765 | -42.992859 | 1,068 |
| Teresópolis | RJT44- | -22.306803 | -42.992922 | 1,069 |
| Teresópolis | RJT46 | -22.306775 | -42.992935 | 1,071 |
| Teresópolis | RJT47 | -22.306783 | -42.992990 | 1,048 |
| Itatiaia | RJII56 | -22.364534 | -44.724839 | 2,269 |
| Itatiaia | RJI57 | -22.364535 | -44.724709 | 2,274 |
| Itatiaia | RJI59 | -22.364497 | -44.724625 | 2,286 |
| Itatiaia | RJI62 | -22.356498 | -44.736398 | 2,151 |
| Itatiaia | RJI64 | -22.356435 | -44.736335 | 2,149 |
| Itatiaia | RJI65 | -22.356403 | -44.736274 | 2,151 |
| Itatiaia | RJI67 | -22.356515 | -44.736454 | 2,151 |
| Itatiaia | RJI68 | -22.356523 | -44.736534 | 2,148 |
| Itatiaia | RJI69 | -22.356506 | -44.736557 | 2,148 |
| Itatiaia | RJI72 | -22.366059 | -44.746456 | 1,951 |
| Itatiaia | RJI74 | -22.366139 | -44.746768 | 1,943 |
| Itatiaia | RJI81 | -22.375046 | -44.755165 | 1,796 |
| Itatiaia | RJI84 | -22.375060 | -44.755070 | 1,794 |
| Itatiaia | RJI86 | -22.375047 | -44.755181 | 1,796 |
| Itatiaia | RJI88 | -22.375020 | -44.755236 | 1,795 |
| Itatiaia | RJI92 | -22.376749 | -44.760301 | 1,669 |
| Itatiaia | RJI93 | -22.376746 | -44.760343 | 1,668 |
| Itatiaia | RJI94 | -22.376723 | -44.760379 | 1,669 |
| Itatiaia | RJI96 | -22.376531 | -44.760754 | 1,672 |
| Itatiaia | RJI97 | -22.376457 | -44.760717 | 1,671 |
| Itatiaia | RJI100 | -22.376572 | -44.760556 | 1,669 |
