## Supplementary figures and images for "Insights from Genetic Studies: SNP Analyses Confirm White Clover Naturalization in Brazil"

### Supplementary figure S1

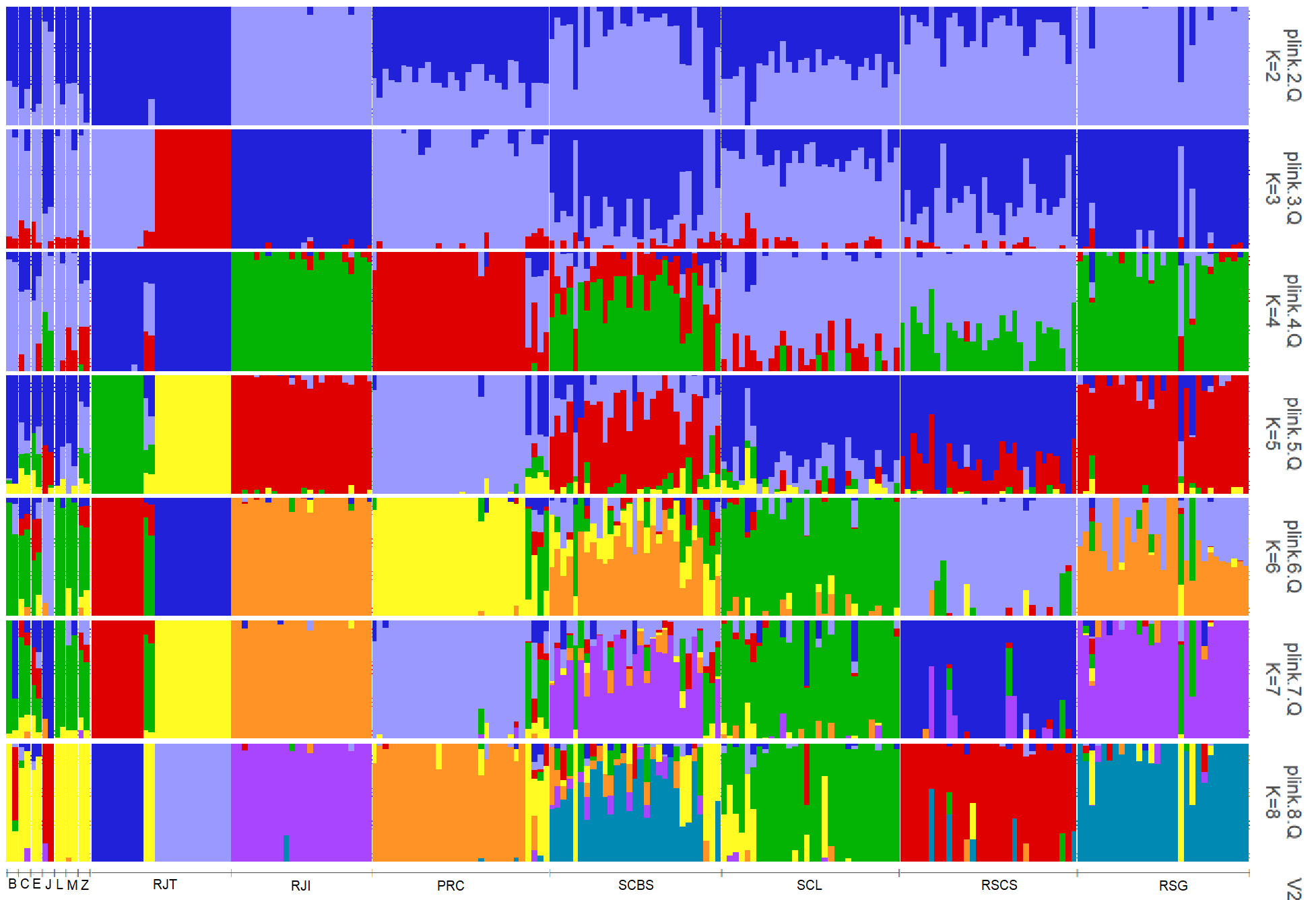
